# Cue integration of texture and elasticity induces roughness metamers in touch

**DOI:** 10.1101/2025.06.02.657485

**Authors:** Karina Kirk Driller, Camille Fradet, Vincent Hayward, Jess Hartcher-O’Brien

**Affiliations:** Delft University of Technology, Landbergstraat 15 2628 CE Delft; Sorbonne University, Institut des Systèmes Intelligents et de Robotique (ISIR), 4 Place Jussieu, 75005 Paris, France; Institute of Neuroscience (IONS), Université catholique de Louvain, Avenue E. Mounier 53, 1200 Brussels, Belgium

**Keywords:** Haptic perception, Texture and material, Cue integration, Touch, Perceptual metamers

## Abstract

Roughness perception is a fundamental dimension of touch that guides object recognition and manipulation. While perceived roughness is typically attributed to surface texture, realworld materials rarely vary in texture alone–they also differ in material properties such as elasticity. Whether material properties contribute to roughness perception, and how they might interact with surface cues, remains poorly understood. Here, we investigated how texture and elasticity jointly influence perceived roughness by parametrically varying both features within a Bayesian optimization discrimination task. Participants compared pairs of stimuli differing in stochastic surface roughness and material elasticity, under both direct and tool-mediated touch. This approach enabled us to estimate two-dimensional perceptual functions and identify haptic roughness metamers–physically distinct stimuli perceived as equally rough. These perceptual equivalences were mirrored in confidence ratings and varied systematically with the relative stiffness between the stimulus and the probing tool or finger, implicating contact-induced vibrations as a mediating factor. Our findings reveal how texture and elasticity cues jointly constrain roughness perception, demonstrating that perceived roughness emerges from the integration of multiple stimulus dimensions rather than surface properties alone. These findings offer practical implications for the design of haptic interfaces and prosthetics, where equivalent percepts may be achieved through different combinations of material and texture, and contribute to a broader understanding of cue integration in haptic perception.

**Significance Statement:** Human touch perception must contend with ambiguity from varying tools, materials, and contexts. This study reveals that perceived roughness arises not from surface texture alone, but from the integration of texture and material elasticity–two physical cues that can trade off to produce indistinguishable tactile roughness percepts. These “roughness metamers” emerge even when touch is mediated through a probe, underscoring the role of vibratory cues, but their emergence depends on the relative stiffness between probe and surface. This finding expands how we understand roughness perception and has direct implications for the design of artificial limbs, robotic sensing, and haptic interfaces that aim to recreate natural tactile experiences.

---

**U**nderstanding how we perceive texture and material properties has long been central to haptic perception research. In this context, the *roughness/smoothness* perceptual dimension has been consistently identified as one of the most important ones, mainly probed through multidimensional scaling studies – alongside the dimensions *hardness/softness* and *stickiness/slipperiness* and sometimes *coldness/warmness* (e.g., (1–4)).

Haptic roughness perception is generally considered a complex multidimensional process that can be influenced by various factors and cues (5–8). In manufactured surfaces, the groove width, ridge height, particle diameter, and the spacing of raised dots, have, for instance, all been related to perceived surface roughness (9–12). Perceived roughness has furthermore been suggested to correlate with various physical properties that arise during dynamic interactions with rough surfaces. These include friction (5, 13), the amplitude of vibration weighted with the frequency response of the Pacinian receptors (14, 15), as well as the average rate of change of the tangential force (16). While one-dimensional roughness parameters such as the arithmetic average (R*α*) and root mean square average (Rq or rms) of the profile height deviations from the mean line exist and are frequently used, no singular definition or predictive roughness feature has been identified to date. Yet, there is general agreement that roughness perception is largely influenced by vibratory information (14, 15, 17, 18). While the so-called duplex theory has highlighted the role of both static and dynamic exploration modes in mediating roughness information (19), dynamic exploration and the generation of vibration become increasingly essential for the discrimination of fine textures (17, 20). This is further supported by findings showing a high correlation between roughness perception via a probe and direct finger contact (21, 22).

While multidimensional scaling studies provide hints towards sensory confounds, controlled experimental studies of roughness perception have typically focused on manipulating surface parameters in isolation, with material properties like elasticity rarely being considered as a potential cue to perceived roughness. Furthermore, experimental research has often relied on either simplified stimulus material, like sandpapers and periodic gratings, to maintain experimental control, or on natural textures and materials that vary across so many dimensions, making it hard to isolate the physical properties that serve as features for perceptual reconstruction of surface roughness. In the physical world, changes in surface features co-exist with changes in material properties (such as elasticity). Natural materials like tree bark, animal hides, or certain food products may, for instance, exhibit rougher textures when older, drier, or more weathered, which may be accompanied by an increased stiffness. As we navigate the world, we rarely encounter individual attributes in isolation, and prior experiences and beliefs can shape our perception (23–25). Therefore, examining how attributes interact and conflict with one another becomes critical when predicting or rendering perception from a reduced cue space (26–29).

In addition to their co-occurrence in the natural environment, there is compelling evidence suggesting that both surface and material cues may play a role in determining the perceived roughness of a surface. For instance, studies on fibrillar surfaces show that material stiffness and surface structure can jointly influence perceived texture (30). It is furthermore known that the elasticity of a material affects its measured and perceived friction (31), although the relationship between friction and perceived roughness is complex (2, 5, 16). Additionally, changing a textured body’s elasticity will likely entail changes in the vibratory pattern created during lateral exploration because the sliding dynamics may be directly shaped by bulk material properties and thus physically relate compliance to surface property perception (32).

The aim of the present study was to investigate whether and how smaller-scale surface topography (here stochastic surface roughness) and material elasticity are jointly encoded during direct and indirect haptic exploration to inform perceived surface roughness. To this end, we carried out a series of roughness discrimination experiments using naturalistic stimuli that systematically varied in their elasticity and stochastic roughness (spanning feature changes between 30 and 500 µm). We hypothesized that a joint influence of surface roughness and material elasticity on roughness perception should lead to perceptual regions of cue fusion in which different combinations of these cues would elicit indistinguishable roughness percepts (metamers). Such metameric relationships, where different combinations of physical stimuli converge on identical perceptual experiences, can offer valuable insights into how the brain integrates different sensory inputs to form a coherent percept (33–36). We further hypothesized that if such a relationship exists for roughness discrimination during direct, dynamic touch and is based on vibration cues, it might persist in indirect touch scenarios. A prior study using stimuli of significantly lower elasticity than the human finger (37) (Chapter 4) failed to demonstrate mixed-cue effects in shaping perceived roughness during direct-touch interactions. The current investigation sought to determine whether adjusting elasticity parameters to more closely approximate the human finger pad’s elasticity range, as well as modifying the probe to better match the stiffness of the stimuli, would reveal the hypothesized confounding cue interactions.

We also measured participants’ confidence in their roughness judgments, allowing us to assess whether potential differences in the accessibility of each stimulus cue were evident not only at the perceptual, but also at the metacognitive level.

## 1. Results

### A. Observations on Exploration Strategy

All participants used lateral movements (stroking) in all conditions, despite being permitted to individually select the most natural exploration strategy.

### B. Discrimination Data

Figure 1 shows a histogram of the area under the receiver operator characteristic curve (ROC AUC) scores for each of the three conditions/blocks. The 99% highest-density interval (HDI) is the shortest interval in which 99% of the posterior mass lies, indicating that the “true” ROC AUC lies within the bounds of the HDI with 99% certainty. As can be seen in figure 1, all HDIs lie above 85%, which is generally considered to be a “good” model fit (38).

**Fig. 1.**
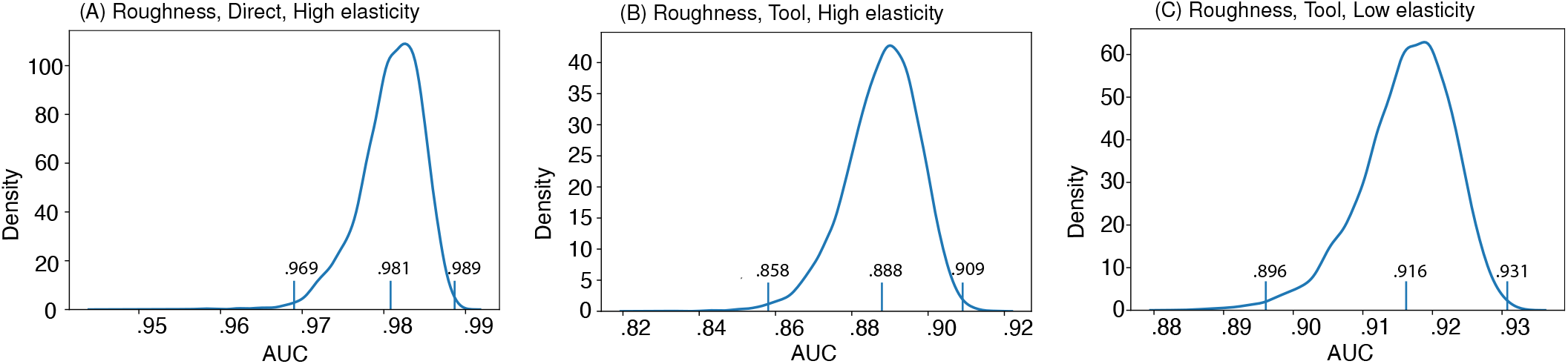
The area under the receiver operator characteristic curve (ROC AUC) of each of the three models. The mean of the distributions, as well as the limits of the 99% highest-density interval (HDI), are labeled.

We then generated psychometric field plots of latent roughness depicting model predictions for our two stimulus parameters (Hurst exponent and Shore value, corresponding to surface roughness and elasticity cues) across all three conditions and individual participants. These visualizations are shown in Figure 2.

**Fig. 2.**
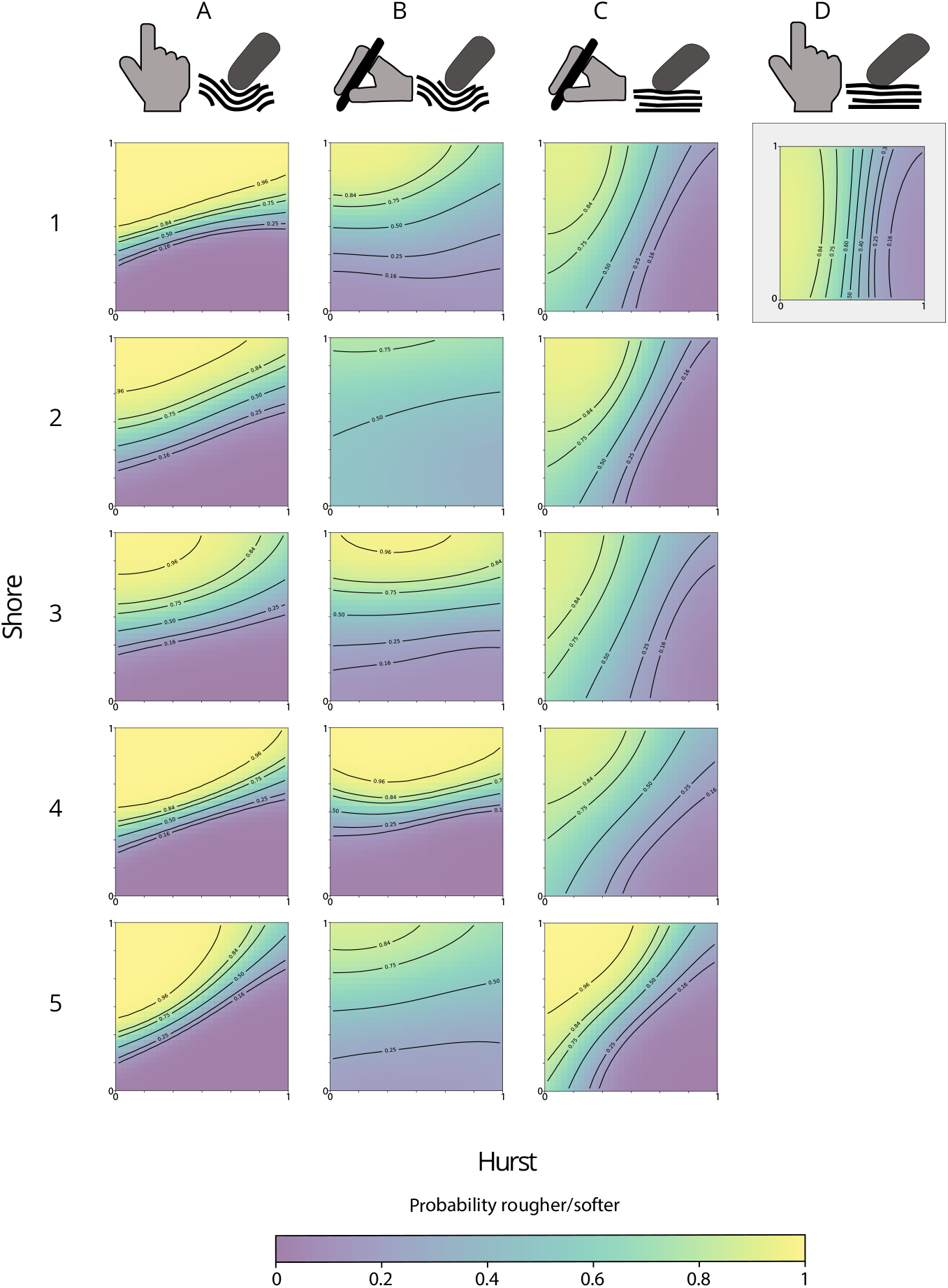
Individual probability plots as a function of Hurst and Shore values. Columns indicate conditions (cf. Table 4), and rows indicate participants. The background color indicates the probability of a stimulus being chosen (as rougher or softer) compared to a stimulus with the medium Hurst and Shore values. Isocontours are plotted at 16%, 25%, 50%, 75%, 84%, and 96%. Because Shore values differ between stimulus sets and are measured on different Shore scales, the plots here correspond to the rescaled [0.1] stimulus space. Min and max values thus correspond to the values indicated in Table 3 for the respective stimulus sets. Note: the Figure in the 4th column for condition D is reproduced from (37) (Chapter 4). It represents the mean probability plot across 15 observers performing the same experiment using direct explorations of the low-elasticity sample set.

The individual probability plots in Figure 2 illustrate the probability that any stimulus within our stimulus space will be identified as rougher compared to a central reference for each subject and condition. The black isocontour lines symbolize the corresponding 16, 25, 50, 75, 84, and 96% probability lines (each isocontour is 1 unit apart in the latent roughness space). Similar to just-noticeable-differences (JNDs), the overall quantity of and the spacing between the isocontours in the space provide a picture of how precisely participants were capable of differentiating within the stimulus space.

Figure 2 reveals remarkably consistent discrimination patterns across participants. A single participant (participant 2) showed a notably reduced discrimination performance in condition B, failing to exceed 75% probability thresholds, suggesting greater difficulty differentiating stimuli in this condition. Nevertheless, the overall pattern remained consistent across participants.

In the individual psychometric fields (Fig. 2), it is clear how roughness discrimination using direct touch of the high-elasticity stimulus set was greatly influenced by the elasticity of the samples, i.e., the Shore value, while it was influenced to a lesser degree by the Hurst exponent (cf. column A). This pattern was largely maintained in roughness discrimination of the same stimulus set using a rigid tool (cf. column B) but with an even greater influence of the Shore elasticity parameter, leaving the influence of the Hurst exponent as almost negligible. It is furthermore visible that the isocontour lines are primarily dictated by the Hurst parameter under this condition. Roughness discrimination of the low-elasticity sample set using a tool, on the other hand, was strongly predicted by both stimulus parameters with a slightly larger influence of the Hurst exponent (cf. column C). These insights were confirmed by examining the best-fitting length-scale values derived from the Gaussian Process (GP) model. These values are displayed in Table 1. The length scales represent the distance along the parameter for which responses are highly correlated and thus indicate the model’s sensitivity to change in the respective parameter. The values have to be viewed in relative terms within each model, where a smaller value indicates a higher sensitivity, and a higher sensitivity indicates a higher weight assigned by the perceptual system in its estimated roughness.

**Table 1.** **Summary of best-fitting lengthscales for Hurst and Shore across the three conditions. Smaller values indicate a greater sensitivity**.

| Condition | A | B | C |
| --- | --- | --- | --- |
| Mode | Direct | Tool | Tool |
| Stimulus Set | High-elast. | High-elast. | Low-elast. |
| Lengthscale Hurst | 1.2047 | 0.7083 | 0.1541 |
| Lengthscale Shore | 0.4655 | 0.0653 | 0.7466 |

### C. Confidence Ratings

Mean confidence ratings were 6.1 (SD = 2.5) for direct roughness discrimination of the high-elasticity sample set (Condition A), 4.69 (SD = 2.56) for roughness discrimination with a tool of the same stimulus set (Condition B), while they were 6.15 (SD = 2.69) for roughness discrimination with a tool of the low-elasticity sample set (Condition C). These results are shown in figure 3.

**Fig. 3.**
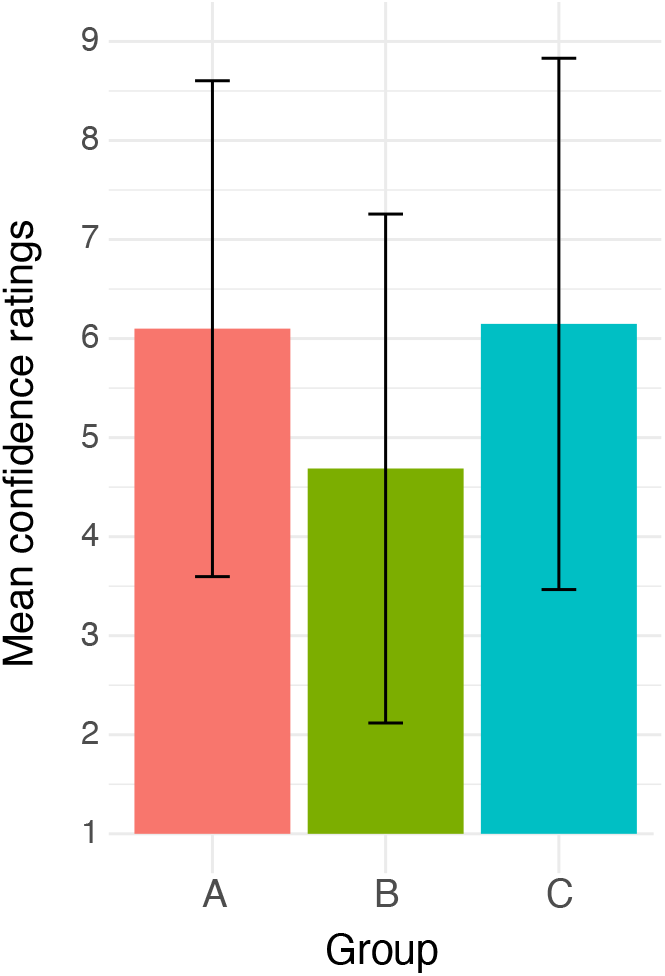
Mean confidence ratings of each of the three conditions of the present study, with error bars showing the standard deviations around the mean.

To assess how stimulus differences influenced confidence ratings, we first calculated standardized differences (ΔHurst and ΔShore) between stimulus pairs by mapping each parameter to a scale of 1–7 at the group level. We then analyzed these relationships using Cumulative Link Mixed Models, with ΔHurst, ΔShore, and their interaction as fixed effects, while accounting for participant-level variability with a random intercept. Perceptual discrimination behavior was highly consistent across participants, supporting the use of group-level modeling. Table 2 summarizes the results.

**Table 2.**
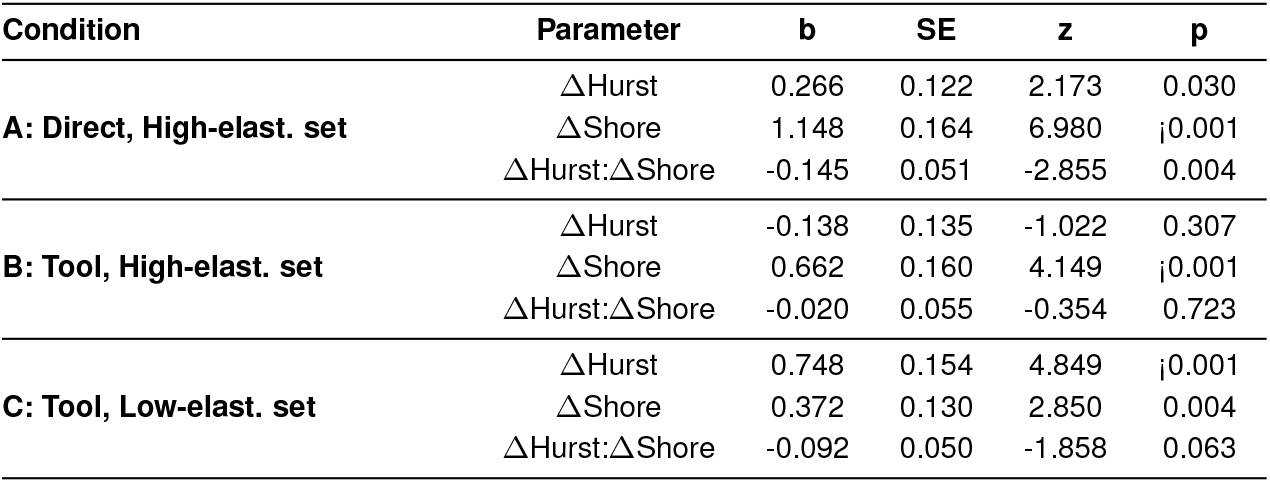
**Results of Cumulative Link Mixed Models for each roughness condition. Coefficients (b) indicate the direction and strength of the relationship between confidence ratings and stimulus differences. The z value represents the standardized effect size; higher absolute values indicate stronger effects. p values denote statistical significance**.

From Table 2, it is evident that ΔShore consistently had a significant positive effect on confidence ratings across all conditions. In Condition A (Direct, High-elast. set), ΔShore had a strong positive effect on confidence (*b* = 1.15, *SE* = 0.16, *z* = 6.98, *p <* .001), as did ΔHurst (*b* = 0.27, *SE* = 0.12, *z* = 2.17, *p* = .030). The interaction was also significant (*b* = *−* 0.15, *SE* = 0.05, *z* = *−*2.86, *p* = .004). In Condition B (Tool, High-elast. set), only ΔShore showed a significant effect (*b* = 0.66, *SE* = 0.16, *z* = 4.15, *p <* .001), while ΔHurst (*p* = .307) and the interaction (*p* = .723) were not significant. For Condition C (Tool, Low-elast. set), both ΔHurst (*b* = 0.75, *SE* = 0.15, *z* = 4.85, *p <* .001) and ΔShore (*b* = 0.37, *SE* = 0.13, *z* = 2.85, *p* = .004) were significant predictors, while the interaction trended toward significance (*b* = −0.09, *SE* = 0.05, *z* = *−*1.86, *p* = .063).

To visualize these differences, we generated predicted confidence landscapes based on our condition dependent CLMM model. These landscapes, which represent the expected confidence ratings for different combinations of Δ Hurst and ΔShore values, can be seen in Figure 4.

**Fig. 4.**
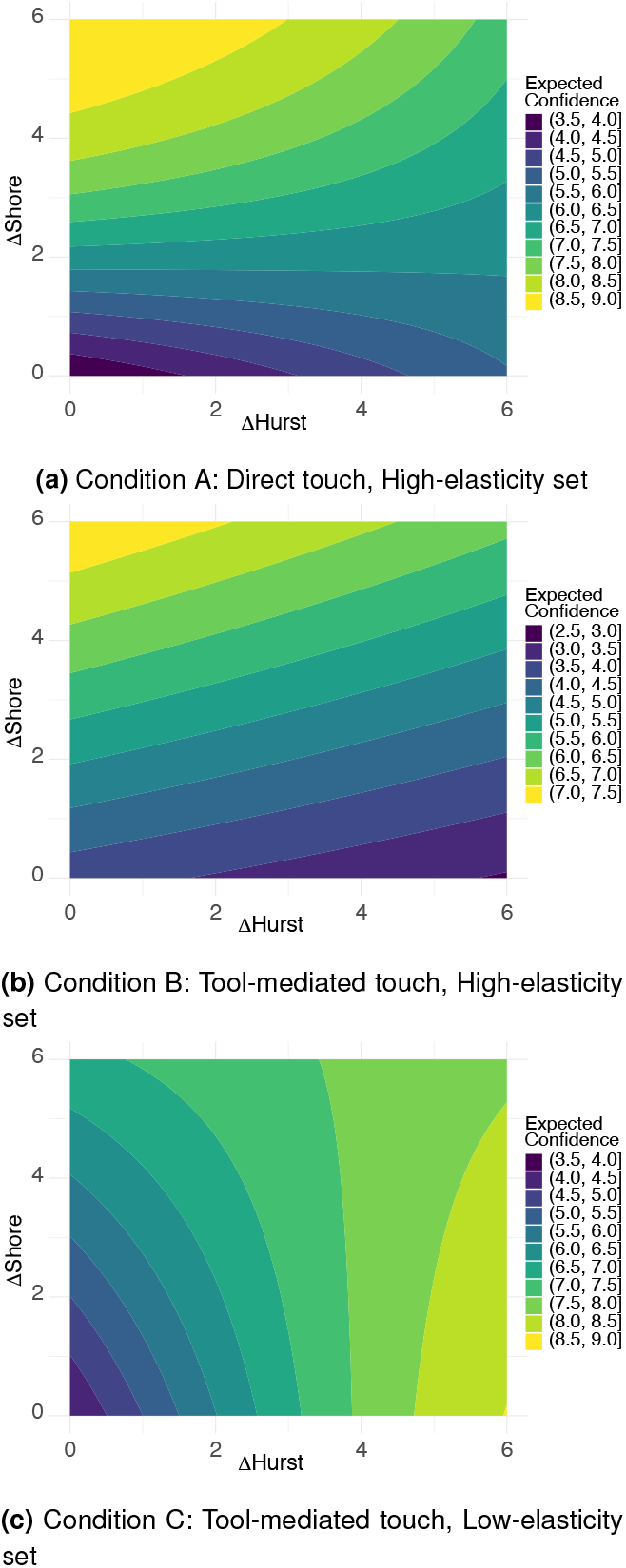
Predicted confidence landscapes based on the CLMM models for each experimental condition for the group. The plots show how the expected confidence ratings vary as a function of stimulus differences in Hurst exponent (ΔHurst) and Shore hardness (ΔShore). Colors represent the predicted confidence.

Figure 4 shows how in condition A, confidence increased systematically with greater stimulus differences, with ΔShore having a notably stronger influence than ΔHurst. The landscape shows confidence levels increasing more steeply along the ΔShore axis, consistent with the higher coefficient for ΔShore compared to ΔHurst (cf., Table 2). A visible interaction effect is apparent, where the influence of ΔHurst diminishes at higher values of ΔShore. For Condition B, the confidence landscape shows an even stronger dominance of the ΔShore axis, aligning with the model results where only ΔShore significantly predicted confidence ratings. The relatively flat gradient along the ΔHurst axis visually confirms participants’ limited sensitivity to Hurst differences when using a tool on the high-elasticity samples. In Condition C, the confidence landscape reveals an influence of both parameters. This is visualized as contour lines that slope more equally with respect to both axes, with slightly stronger effects of ΔHurst overall.

Together, the data in this study support a model in which both surface features and material elasticity contribute to perceived roughness, both for direct and indirect touch. However, this relationship is dependent on the relative range of elasticities tested (high vs. low) and the type of probe used (finger vs. tool), with elasticity sometimes even dominating roughness perception. Combined effects of surface features and elasticity were also observed in confidence ratings during roughness discrimination, with differences in each parameter predicting differences in confidence ratings to varying degrees.

## 2. Discussion

We conducted a series of roughness discrimination experiments to explore the interplay between stochastic surface roughness and material elasticity in shaping roughness perception during both direct and indirect touch interactions. A previous study using lower-elasticity stimuli (37) (chapter 4) did not capture mixed-cue effects, likely due to a mismatch between stimulus and finger elasticity. The present study addressed this by aligning sample elasticity with the human finger pad and adjusting probe properties accordingly, successfully revealing cue confounds. Analyses of confidence ratings ratings indicated that the presence of cue interactions in higher-level metacognitive processing as well. The findings and their implications are discussed in detail below.

### A. A Roughness Metamer

We observed perceptual constancy of roughness across physical stimulus parameter changes, highlighting a clear interdependency between surface features and material elasticity in informing perceived roughness. Across all participants and conditions, roughness discrimination was systematically influenced by both stochastic surface roughness and material elasticity properties. Our data clearly identified metameric regions in the cue space where distinct combinations of elasticity and surface roughness produced perceptually equivalent roughness experiences. The diagonal orientation of isocontour lines in Figure 2 demonstrates that higher surface roughness (lower Hurst exponent) combined with greater elasticity (lower Shore value) produced perceptual equivalence to lower surface roughness paired with lower elasticity. This metameric relationship held for both direct and indirect dynamic touch interactions, indicating that roughness perception integrates information across multiple physical dimensions rather than being determined by surface properties alone. This finding has immediate practical implications for haptic interface design, as it suggests that equivalent roughness sensations can be achieved through strategic combinations of surface and material properties.

### B. Vibration as a Cue: A Propagating Metamer

Roughness perception has long been acknowledged as a complex and multidimensional process (5–8, 12), but the direct effects of changes in the elasticity of a textured surface on its perceived roughness are reported here for the first time. This novel finding demonstrates that roughness perceptual constancy is maintained across various physical parameter changes.

Friction, which can be influenced by both surface texture and material elasticity, could be one possible mediator. Since friction between the fingertip or probe and a rough surface is related to the real contact area between them (e.g., (39, 40)), it is likely that variations in both elasticity and surface features will have had an influence on the friction dynamics in the present study. However, the relationship between friction and both perceived and physical surface roughness is complex (2, 16). On a micro-scale, friction often decreases as surface roughness increases, but this pattern inverts with higher degrees of roughness (5, 41, 42), although greater friction is often associated with greater perceived roughness (1, 5, 43). Consequently, exceptionally smooth or flat surfaces can sometimes be perceived as rougher than textured ones, due to their increased resistance to sliding and the potential for increased large-scale stick-slip (5). High elasticity, on the other hand, is sometimes associated with increased friction (31, 39, 42). For instance, one study found that both measured and perceived friction *increased* as the stiffness of micro-structured rubber samples *decreased* (31). However, we did not observe that more elastic surfaces were perceived as rougher in the present study for any of the interaction types, but the reverse. Although friction may therefore have played a role in the perceived roughness of the samples in the present study, it likely played a secondary role in producing the observed metamers.

It is generally well-established that surface roughness estimates rely heavily on vibratory information (e.g., (19, 44, 45)) and that subjects can discriminate complex textures on the basis of vibration cues alone (46). This raises pertinent questions about the vibratory cues as participants interacted with the surfaces in the present study. Specifically, changing a textured body’s elasticity likely alters tribological events during lateral exploration such as ploughing, adhesion, and impact frequency responses, affecting the resulting vibration (47, 48). Thus, a mechanistic interpretation may suggest that more elastic samples deform more upon dynamic touch (i.e., ploughing events), resulting in decreased impact and indentation on the finger/probe during lateral exploration. Magnitude and frequency of the resulting vibrations during sliding may consequently directly depend on the bulk elasticity (32).

The impact of vibratory cues on the final perceptual outcome of a roughness judgment becomes particularly evident in tool-mediated interactions. During indirect touch, vibratory cues are preserved but transformed by the tool’s mechanical properties as they propagate to the hand (45, 49, 50). In the present study, roughness discrimination of the more elastic stimulus set was remarkably similar for direct and indirect touch, although a shift towards a stronger influence of the Shore value emerged during indirect touch, probably due to the greater impedence mismatch between the tool and the sample (see section The Importance of Relative Stiffness). However, the persistence of this metameric relationship across both interaction modes–whether local skin input was available or not–underscores the primacy of vibratory information in shaping the final percept. It suggests that participants did not selectively subtract or compensate for the elasticity cues in their roughness judgments, even when local tactile information was directly available through finger contact. Instead, elasticity appeared to be embedded in the vibratory signal itself and not easily separable from surface texture during perceptual processing.

### C. The Importance of Relative Stiffness

Our findings demonstrate that cue confounds in roughness perception—where surface texture and material elasticity jointly shape perceptual judgments—emerge reliably when the stimulus elasticity falls within a behaviorally relevant range relative to the probe. This pattern was observed for both direct and indirect touch with the high-elasticity sample set, and for indirect touch with the low-elasticity set. These results highlight that relative, not absolute, stiffness between the probe and the sample is critical. When a stimulus is much stiffer than the finger or probe, changes in its elasticity exert minimal influence on perceived roughness. However, when the stimulus elasticity is comparable to or lower than that of the probe–whether it is the finger or a rigid tool–material elasticity can significantly influence roughness perception, at times even more than surface texture (see Fig. 2, Table 1).

When probe and sample exhibit similar resistance to movement and deformation, the two surfaces can be said to be impedance matched (51–53), a relationship grounded in the concept of the equivalent modulus for two elastic bodies in contact (54, 55). Although contact mechanics were not directly measured here, our data support the interpretation that cue integration effects are strongest when the stiffness of the probe and sample are closely matched. In these cases, changes in sample elasticity substantially shape the frequency response of sliding interactions and thus the vibratory input driving roughness perception. Figure 5 illustrates this relationship through four representative scenarios.

**Fig. 5.**
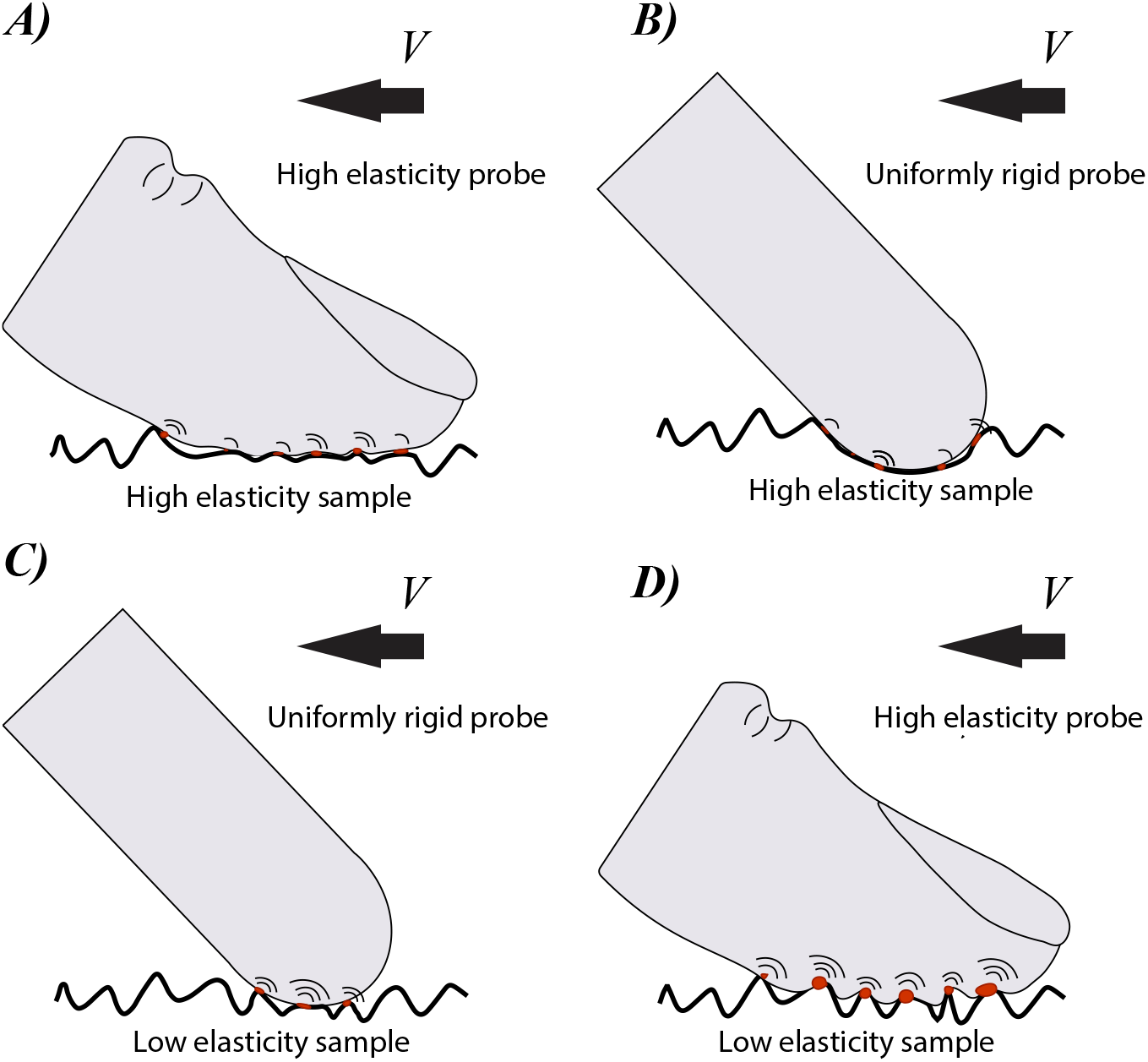
Four scenarios illustrating the effect of the relative elasticity of a textured sample and the probe used during sliding in the present study. **A)** High-elasticity probe and sample: both deform; vibration reflects a combined influence of elasticity and surface texture. **B)** Rigid probe and high-elasticity sample: sample deforms easily; elasticity dominates, texture plays a reduced role. **C)** Rigid probe and low-elasticity sample: sample partially resists deformation; asperities push back against the probe, enhancing ploughing and contact forces. Joint effects of elasticity and texture. **D)** High-elasticity probe and low-elasticity sample: sample is significantly stiffer than probe and largely resists deformation; variations in elasticity within this range have minimal impact. Surface texture dominates the vibratory signal.

**Fig. 6.**
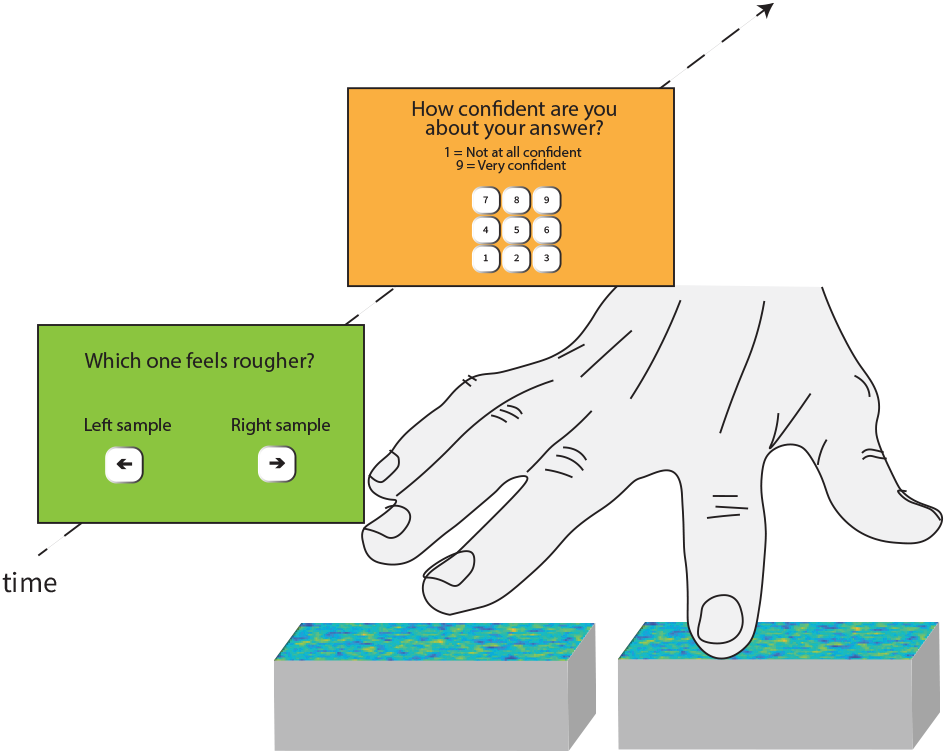
Procedure

Within a given range of relative sample elasticity to the probe, changes in the elasticity of the sample should significantly alter the frequency response of dynamic impacts during lateral exploration. In Figure 5, scenario A, the probe and sample deform against each other. Changes in the material elasticity within this range should change both the dynamics of adhesion, ploughing, and the resulting frequency response of impacts. However, if the probe is significantly more rigid than the sample (Fig. 5, scenario B), sample asperities will deform and oppose little to no resistance to the probe. Here, the actual shape of the asperities may thus have little impact on the frequency response and the resulting roughness percept. In the scenario of a uniformly rigid probe and a sample low in elasticity (Fig. 5 scenario C), sample asperities will deform but still oppose a significant resistance to the probe. Changes in the elasticity of the sample within this range should result in changes in ploughing, adhesion, as well as the resulting frequency response of impact events propagating through the probe. Finally, if a textured surface and its asperities are significantly more rigid than the probe (Fig. 5, scenario D), little or no deformation of the sample should take place during dynamic interactions, and impact events will be absorbed by the probe’s material elasticity which deforms around the sample. Since the real-contact area is unlikely to change with changes in the elasticity of the sample, no significant changes in adhesion should be present either. Within this range, the resulting vibration and roughness percept should be predominantly defined by surface features.

Together, these scenarios illustrate how the relative elasticity of the probe and sample determines the frequency content of vibrations arising during dynamic touch. As demonstrated in this study, these interactions can give rise to cue confounds when material and surface properties jointly shape roughness perception.

For haptic technology applications, this implies that interface compliance becomes an active determinant of perceptual experience. Touch-based interfaces could thus be designed to exploit optimal stiffness matching to enhance texture discrimination or achieve desired perceptual outcomes through material selection, without requiring complex surface actuation.

### D. Metameric Cue Confounds and Confidence

Across conditions, confidence ratings reflected the relative accessibility and informativeness of the available cues during roughness discrimination at the group level. Confidence was highest when both stimulus parameters—surface texture and elasticity—contributed meaningfully to the task (conditions A and C), and slightly lower when one cue dominated while the other became less informative (condition B).

Specifically, in Condition A (direct touch, high-elasticity set), confidence increased with greater differences in both Shore and Hurst values, suggesting that both cues were accessible and behaviorally relevant. In Condition C (tool touch, low-elasticity set), a similar pattern emerged—though with a slightly stronger influence of Hurst—indicating that cue integration was preserved even without direct skin contact. In contrast, in Condition B (tool touch, high-elasticity set), confidence was overall lower and more strongly driven by Shore differences, pointing to a reduced availability of texture-based information. This aligns with the idea that the stiff probe in this condition limited the transmission of vibratory cues from surface features (see Section The Importance of Relative Stiffness).

Taken together, these results suggest that confidence ratings may reflect the clarity or diagnostic value of the integrated sensory input, rather than the mere presence or number of cues. While all conditions involved cue fusion, the effectiveness of that fusion—and thus the certainty it afforded—appeared to vary with the probe–surface interaction. While these results are preliminary due to the small sample and group-level analysis, the findings tentatively support the view that metacognitive access is shaped by how reliably cues support perceptual decisions, rather than by whether cues are fused or separated. This perspective contrasts with interpretations that cue conflicts or ambiguities necessarily reduce confidence (e.g., (56)) and is more consistent with the idea that cue fusion can be metacognitively seamless when cues are compatible and behaviorally useful. Future work should examine this hypothesis more directly, ideally with individual-level modeling and larger samples.

## 3. Conclusions

We investigated the combined influence of surface features (stochastic roughness) and material elasticity on perceived roughness during both direct and tool-mediated touch using two different stimulus sets. We demonstrated how material elasticity can serve as a behaviorally relevant cue to roughness perception depending on the relative elasticity of the stimuli and a rigid probe. In doing so, we identified a perceptual roughness metamer, where different combinations of material elasticity and surface roughness can result in the same subjective roughness. This metameric relationship was observed in both direct and indirect touch conditions, but the relative contribution of each cue varied systematically with the stiffness match between stimulus and probe (i.e., rigid probe or finger). These findings challenge models that attribute roughness perception solely to surface parameters, instead underscoring its multidimensional nature–integrating both surface and material properties. A critical implication of this finding is greater flexibility in conveying roughness in haptic applications. Subjective roughness can be modulated not only via changes in stimulus texture or material but also through adjustments to the probe or interface itself. For example, the stiffness and composition of an artificial limb will directly influence how surfaces and textures are perceived through it.

## 4. Materials and Methods

### A. Participants

Five healthy adult volunteers (2 women, 3 men), average age of 32.2 (SD 2.05) participated in the study. Small sample studies like this are customary in sensory psychophysical research where the focus is on measurement reliability and repetitions within subjects, and the experimental power is concentrated at the individual participant level (57, 58). All participants reported being right-handed and carried out the experiment with their right hand. All participants gave written informed consent prior to the study. The methods of this study were performed in accordance with relevant guidelines at Sorbonne University and the Declaration of Helsinki. No participants or trials were excluded after data collection.

### B. Apparatus

Participants were seated in a chair at a desk in front of a computer screen. During the experiment, they wore noise protection headphones to ensure that the feedback they received was purely haptic in nature. The light in the experimental room was dimmed so that differences between the stimuli could not be seen, while the outline of the stimuli could still be made out for targeted interactions. Stimuli were placed in front of the participant’s dominant hand for semi-controlled exploration with the index finger or a rigid tool (the rounded backside of a wooden paintbrush). A numberpad was provided for responses via keypress at the side of the participant’s non-dominant hand.

### C. Stimuli

Two stimulus sets were used in the present study. First, we made use of the 3D-printed database of 49 stochastically rough, self-affine stimuli, systematically varying in their stochastic surface roughness (Hurst exponent) and material elasticity as reported in detail here (59). The surface variations in these stimuli spanned feature changes between 30 and 500 µm, bridging both micro-, meso-, and macroscale features relative to a fingertip. This stimulus set was also used in a previous experiment for direct-touch interactions (37). Elasticity and roughness parameters of this database are summarized in Table 3 A and B, while a precise characterization of it can be found here (59).

**Table 3.** **Roughness parameters (A) of both stimulus databases and elasticity parameters of the original low-elasticity stimulus database (B) and the new high-elasticity database (C). N.B. different Shore scales in tables B and C**

| A: Surface roughness parameters of both stimulus sets |  |  |  |  |  |  |  |
| --- | --- | --- | --- | --- | --- | --- | --- |
| Surface Nr. | S1 | S2 | S3 | S4 | S5 | S6 | S7 |
| Hurst exponent | 0.3 | 0.4 | 0.5 | 0.6 | 0.7 | 0.8 | 0.9 |
| Rq [$\times 10^{-2}$ mm] | 5.68 | 5.16 | 4.79 | 4.51 | 4.30 | 4.14 | 4.01 |

**B: Elasticity parameters of the low-elasticity sample set taken from (59)**
| Elasticity Nr. | E1 | E2 | E3 | E4 | E5 | E6 | E7 |
| --- | --- | --- | --- | --- | --- | --- | --- |
| Shore-A value | 24 | 25 | 26 | 29 | 34 | 44 | 66 |
| Y. Modulus (MPa) | 0.121 | 0.122 | 0.128 | 0.144 | 0.179 | 0.264 | 0.611 |

**C: Elasticity parameters of the high-elasticity stimulus set created here**
| Elasticity Nr. | E1 | E2 | E3 | E4 | E5 | E6 | E7 |
| --- | --- | --- | --- | --- | --- | --- | --- |
| Shore-00 value | 9 | 18 | 25 | 32 | 39 | 43 | 47 |
| Y. Modulus (MPa) | 0.017 | 0.029 | 0.041 | 0.055 | 0.072 | 0.084 | 0.097 |

In addition, we created a new stimulus set with the same surface parameters as the original database, but with an updated elasticity range close to the elasticity of a human finger pad. This was done by creating negative molds using Ecoflex™ 00-30 silicone. While resulting in an inversion of peaks and valleys, the statistical roughness of these stimuli could be considered the same as that of the previous stimulus set.

Since most commercially available stereolithography (SLA) resins inhibit the curing of polydimethylsiloxane (PDMS), we first printed an additional set of 7 rigid surface samples suitable to be used as molds for the final silicone specimens. To avoid curing inhibition, we followed the steps outlined by Venzac et al. (60) in the fabrication of these surfaces. The rigid surface samples were printed using a Formlabs Form 3 Printer and grey resin, which has a similar resolution to the Polyjet 735 used for the printing of the original stimulus database. After printing, the samples were washed in IPA for thirty minutes in a Formlabs Form Wash station and carefully rinsed manually with isopropanol using a soft paint brush. The samples were then cured for twenty-four hours at 60°C. Each of these seven rigid surfaces was then mounted into

3D-printed frames, together creating molds for the uncured silicone mass. These frames were printed in PETG and covered in Kapton tape to further prevent potential curing inhibition at the sides of the molds. Upon mixing the two parts of the silicone, a total mass of 18 g was carefully poured into each of the molds and then removed again after curing. To achieve different elasticities, we adjusted the mixing ratio of the two-part elastomer Ecoflex™ 00-30. The above procedure was therefore repeated seven times, with an adjusted mixing ratio for all the 7 surfaces. This resulted in a final database of 49 silicone rubber samples sized 50x31x13 mm. Their Shore hardness was subsequently measured using a Shore-00 durometer (ASTM 2240) and the Young’s modulus was calculated using Gent’s equation (61). Table 3C summarizes the achieved elasticity values for this stimulus database. Detailed information about the precise mixing ratios used to achieve the different elasticities can be found in SI Appendix 1.

After fabrication and before every data collection, all specimens were coated in talcum powder to reduce adhesion differences caused by varying elasticity, since softer materials are universally more adhesive (62). A pilot study had ensured that elasticity and surface cues were discernible within the cue space using both stroking and pressing in the absence of the other cue.

### D. Experimental Design and Procedure

We used an adaptive exploration procedure based on the non-parametric Bayesian inference framework Aepsych (63, 64)) to model a single latent function *F*:

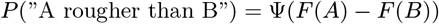

determining the probability of the perceptual judgment between any two stimuli in the stimulus space. The precise configuration of the model is specified in the SI Appendix appendix 2.

Participants carried out a 2AFC-discrimination task, in which they had to indicate which one of two stimuli felt rougher. Each trial began with a window appearing on the participant screen indicating “Which stimulus feels rougher?”. Participants were instructed to use free natural explorations of the surfaces, using any interaction methods they wanted (65), but only using their dominant index finger in the direct-touch condition or the provided tool in the indirect-touch condition. They were asked to explore the left stimulus first and then move to the right stimulus. They were also asked not to explore the edges and corners of the samples and to avoid using their fingernails. Participants were encouraged to give quick and intuitive answers. After their response, participants were asked to rate how confident they were about the answer given on a Likert scale from 1-9. All answers were given by keypress with the non-dominant hand. After each trial, the experimenter placed a new pair of samples in front of the participant, whereafter the task window reappeared on the participant’s screen, indicating that they could commence with the next trial.

The experiment consisted of three conditions, each constituting a block comprising 50 trials. Each block was interspersed with a 5–10-minute break. The approximate duration for each block was 30 minutes. The blocks were designed to cover three different combinations of *exploration mode* (direct/tool), and *stimulus set* (high-/low-elasticity). Note, direct explorations of the low-elasticiy sample set were explored in a preveous study and were therefore not included here. Table 4 summarizes the three conditions.

**Table 4.** The three conditions included in this study, each constituting one experimental block.

| Condition | Mode | Stimulus set |
| --- | --- | --- |
| A | Direct | High-elastic. set |
| B | Tool | High-elastic. set |
| C | Tool | Low-elastic. set |

### E. Statistical Analyses

A Gaussian Process (GP) model (66) was fit to each of the three conditions. The model used Houlsby’s (67) pairwise kernel for the Hurst and Shore values and an index kernel for participants. The model posterior was estimated using variational inference (68), with a hyperparameters fit using maximum likelihood estimation. The Hurst and Shore values (cf. the corresponding values in Table 3 in the methods section) were min-max scaled to the range [0, 1]. Formally, given the binary nature of the outcome, we assume each roughness judgment, *y*, to be drawn from a Bernoulli distribution with probability Φ(*f* (*u, v, p*)), where Φ is the Gaussian Cumulative Distribution Function (CDF), and *f* (*u, v, p*) is a latent function over the model inputs: the left and right stimuli (given by their Hurst and Shore values) and participant index.

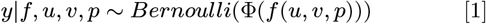

We placed a Gaussian Process (GP) prior on *f* . As is typical in GP regression, we used a constant mean function of zero, and we used a covariance function that is the product of two different covariance functions, *k*_stim_ and *k*_part_.

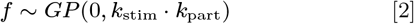

The function *k*_stim_ models the covariance between pairs of stimuli. Given two pairs [*a, b*] and [*c, d*], the covariance is given by

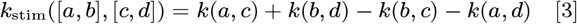

where *k* is the Matérn 5*/*2 kernel (the derivation for this expression is provided by Houlsby et al. (67)). The function *k*_part_ is an index kernel, which models the covariance across participants. The GP approach provides a single model that considers the combined impact of hurst exponent and elasticity on perceived roughness, while also accounting for individual observer differences and cross-observer correlations.

Model fits were subsequently assessed using the area under the receiver operator characteristic curve (ROC AUC) (38), which measures how well the model is able to predict which of two stimuli will be judged as rougher. To estimate the distribution of predictions, 10,000 samples were drawn and the ROC AUC was calculated for each of those samples for each model.

We then created plots of the model predictions for each of the three conditions. These predictions were made on a dense grid corresponding to our two stimulus parameters (Hurst and Shore).

Confidence ratings were first compared using means and standard deviations. Likert-scale data are technically considered ordinal, but can in some cases be considered approximately continuous (especially in the context of a larger numerical range) (69). We report means, rather than medians or modes, as they have the advantage of being easy to locate on the original scale. However, Generalized Linear Mixed Models (ordinal package, “clmm” function in R (70)) were used to evaluate the effects and interactions of changes in Hurst and Shore values on the confidence ratings.

To evaluate the relationship between a participant’s confidence ratings and changes in the stimulus parameters within a trial, ΔHurst and ΔShore values were calculated by first standardizing each parameter range to values ranging from 1 to 7, and then computing the absolute difference between the values of each stimulus pair within a trial. This was done separately for each condition. Cumulative Link Mixed Models (CLMMs) were then fitted to the data for each condition using ordinal logistic regression. The models included ΔHurst, ΔShore, and their interaction as fixed effects, with a random intercept for participants to account for individual variability. Model fitting was performed using the Laplace approximation.

To visualize the differences between conditions, we generated predicted confidence landscapes based on the CLMMs, representing the expected confidence ratings for different combinations of ΔHurst and ΔShore values. A group-level modeling approach was chosen because individual-level modeling was not feasible given the ordinal nature of the confidence ratings and the limited number of trials per condition.

Statistical analyses of the discrimination data were conducted using Python (Anaconda Navigator, Spyder version 5.4.3) and using AEPsych (63, 64). Analyses of the confidence ratings were performed using R (version 4.3.1, R Core Team, 2021).

## ACKNOWLEDGMENTS

We thank Craig Sanders for his valuable guidance on the data analysis.

