## supplementary material for "Cue integration of texture and elasticity induces roughness metamers in touch"

**This PDF file includes:**

Supporting text
Fig. S1
Table S1
SI References

### Supporting Information Text

### S1. Ecoflex™ Mixing Ratios

Ecoflex™ 00-30 is a two-part silicone elastomer, where parts A and B are mixed in equal proportions (a 1:1 ratio) to achieve a Shore-00 hardness of approximately 30. The Shore-00 scale measures the material's hardness, with higher values signifying increased hardness. Modifying the ratio of parts A to B allows for the tuning of the elastomer's hardness or elasticity. Typically, decreasing the amount of part B relative to part A leads to a harder material with a lower elasticity and higher shore value (e.g., (1)). Table S1 documents the mixing ratios used for the stimuli created, indicating the quantity of part B for every one part of A.

It is salient from Table S1 that the relationship between the proportions of parts A and B, where a greater proportion of part A typically results in a higher Shore 00 value, inverts beyond a certain threshold. Specifically, when the amount of part B is significantly reduced relative to part A, the trend reverses, and a further decrease in part B results in specimens with a lower Shore value. Samples E1-E3 demonstrate this behavior. This phenomenon is further illustrated in Figure S1.

**Table S1. Silicone mixing ratio**

| **Elasticity Nr.** | E1 | E2 | E3 | E4 | E5 | E6 | E7 |
|---|---|---|---|---|---|---|---|
| **Ratio part B per 1A** | 0.125 | 0.146 | 0.16 | 2 | 1 | 0.5 | 0.25 |
| **Shore-00 value** | 9 | 18 | 25 | 32 | 39 | 43 | 47 |

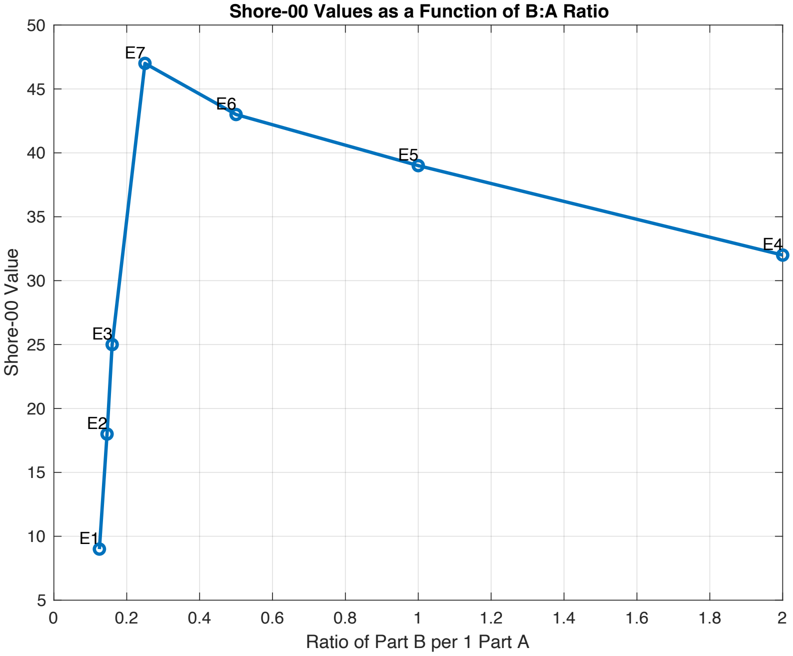


**Fig. S1.** Achieved Shore-00 hardness as function of Ecoflex™ 00-30 mixing ratio for the stimuli

### S2. AEPsych Model Configuration

AEpsych uses state of the art machine learning models but is designed to allow users to configure experiments using 'config files' without directly interacting with the underlying code. The config file used in the experiments is presented below:

```
config = {
'[common]',
'parnames = [Hurst, Shore]',
'lb = [1, 1]',
'ub = [7, 7]',
'outcome_type = pairwise_probit',
'strategy_names = [init_strat, opt_strat]',
'[init_strat]',
'n_trials = 40',
```

```
'generator = PairwiseSobolGenerator',
'[opt_strat]',
'n_trials = ' num2str(n_trials_opt_strat) '',
'refit_every = 5',
'generator = PairwiseOptimizeAcqfGenerator',
'acqf = PairwiseMCPosteriorVariance',
'model = PairwiseProbitModel',
'[PairwiseMCPosteriorVariance]',
'objective = ProbitObjective',
'[PairwiseProbitModel]',
'inducing_size = 100',
'mean_covar_factory = default_mean_covar_factory',
'[PairwiseOptimizeAcqfGenerator]',
'restarts = 10',
'samps = 1000',
};
```

While the sobol or initiation trials were thus set in the config, the total number of trials (50) was configured in the experimental matlab code. Furthermore, because the AEPsych server provided sampling parameters between 1 and 7 (a Hurst and a Shore value) that were continuous, these values were rounded up or down to the closest whole numbers, corresponding to the discrete number of stimuli (49) that were available in our 2D-stimulus space. In order to avoid that the same physical stimulus would be selected twice, which would have required a duplicate of each stimulus, each point in the server space was matched to corresponding points in the sample space based on euclidean distances between the continuous space points and the discrete space points.
